# Cth1 is essential for gametogenesis and fertility in zebrafish

**DOI:** 10.64898/2026.08.26.747083

**Authors:** Gopal Kushawah, Stephanie H. Nowotarski, Carmichael Carrie, Malloy Seth, Hannah Wilson, Ning Zhang, Ariel A. Bazzini

**Affiliations:** Stowers Institute for Medical Research, 1000 E 50th Street, Kansas City, MO 64110, USA; Department of Molecular and Integrative Physiology, University of Kansas Medical Center, 3901 Rainbow Blvd, Kansas City, KS 66160, USA

**Keywords:** CRISPR-Cas9, *Cth1*, Embryonic lethal, Germ cells, Gametogenesis, RNA-decay factor, Zebrafish

## Abstract

Infertility frequently arises from defects in germ cells and early embryonic transition. Successful fertilization depends on the developmental competence and molecular integrity of mature gametes from both parents, which are established through tightly coordinated programs of RNA regulation, metabolism, and genome maintenance. In our previous work, we identified Cth1 as a maternally regulated RNA-decay factor essential for early embryonic development, acting through spatiotemporal control of maternal transcript clearance. Interestingly, *Cth1* loss of function in adults also resulted in infertility, suggesting an additional and unexplored role during gametogenesis. Here, we extend these findings by defining the gametogenic function of Cth1 in zebrafish. Through detailed phenotypic, cytological and molecular characterization of Cth1 loss-of-function mutants, we show that Cth1 is highly enriched in germ cells and early embryos and is spatio-temporally localized across oogenesis and early development. Loss of Cth1 causes severe defects in early oogenesis and spermatogenesis, resulting in complete infertility in males and females. Mutant germ cells display transcriptomic changes consistent with metabolic and translational dysregulation, increased DNA damage, and striking abnormalities in gamete morphology. Together, these findings identify Cth1 as essential for gamete quality and fertility. Our study suggests a link between RNA decay–mediated regulation of metabolism and genome integrity during germ cell development and reveals disruption of post-transcriptional control as a potential mechanism underlying infertility.

**Highlights:** xxxx

## Introduction

Infertility represents a major global health burden, affecting an estimated 10–15% of couples worldwide (Liu et al, 2025; Vander Borght & Wyns, 2018), and frequently arises from defects in germ cell development, gamete quality, and early embryonic transitions (Carson & Kallen, 2021; Vander Borght & Wyns, 2018). The ability of mature gametes to support fertilization and development depends on tightly coordinated gene-regulatory programs that preserve germ cell identity, ensure meiotic progression, and maintain cellular and genomic integrity (Belloc & Mendez, 2008; Gao et al, 2024; Innocenti et al, 2022). However, during key phases of gametogenesis and early embryonic development, transcription is limited, silenced, or uncoupled from protein synthesis (Bellutti et al, 2025; Guo et al, 2022; Xu et al, 2012). Consequently, post-transcriptional regulation becomes a dominant mode of gene control in the germline, governing mRNA stability, localization, poly(A) tail dynamics, and translational timing (Schultz et al, 2018; Sha et al, 2020a; Sha et al, 2020b; Tadros & Lipshitz, 2009; Vastenhouw et al, 2019). RNA-binding proteins (RBPs), including DAZL (Deleted in Azoospermia-Like), Ddx4/VASA, Nanos, PUMILIO, CPEB-family proteins, and MSY2/YBx2, are central to these processes, acting as critical regulators of developmental competence and cellular homeostasis (Forbes & Lehmann, 1998; Ivshina et al, 2014; Jiang et al, 2025; Medvedev et al, 2011; Tsuda et al, 2003; Xu et al, 2021; Yang et al, 2020). Disruption of RBP-mediated regulatory networks has emerged as a recurring molecular feature underlying infertility, germ cell quality, and early developmental failure (Chukrallah et al, 2023; Idler & Yan, 2012; Peart et al, 2022; Vong et al, 2021; Zagore et al, 2015).

1. *Cth1* is a highly abundant mRNA in zebrafish germ cells and early embryos.
2. Loss of *cth1* causes complete infertility in both male and female zebrafish due to gametogenic failure.
3. *Cth1* mutant gametes exhibit metabolic and translational dysregulation, chromatin instability, and elevated DNA damage.
4. Cth1 is essential for oocyte growth and maturation, and for sperm flagellar architecture and quality control.

The ZFP36/TTP family of CCCH zinc-finger RNA-binding proteins functions as critical post-transcriptional regulators of RNA stability (Cook et al, 2022; Wells et al, 2017). In vertebrates, ZFP36 family members, such as ZFP36l2 and C3H-4 promote RNA turnover by deadenylating transcripts with AU-rich elements within 3′ untranslated regions and recruiting deadenylation and decay machinery, including the CCR4–NOT complex (Belloc & Mendez, 2008; Wells et al, 2017). Although these proteins are best known for regulating transcripts associated with inflammation, stress-response, proliferation, and differentiation in somatic tissues, emerging evidence suggests that related RNA-decay pathways may also contribute to reproductive competence (Cook et al, 2022; Snyder et al, 2024). In *C. elegans*, the CCCH-type zinc-finger protein OMA-1 is known to regulate oocyte maturation and early meiotic progression (Kaymak & Ryder, 2013; Shimada et al, 2002). In Xenopus oocytes, the C3H-4 protein is known to regulate deadenylation during different phases of meiosis and loss of C3H-4 induces meiotic arrest during first meiotic division (Belloc & Mendez, 2008). In mice, ZFP36l2 is required for female fertility (Ball et al, 2014; Ramos et al, 2004). In humans, biallelic variants in ZFP36L2 have been associated with female infertility, characterized by oocyte maturation defects or recurrent preimplantation embryonic arrest (Wan et al, 2024; Zheng et al, 2022; Zhou et al, 2023). These studies support an important role for ZFP36-family RNA-binding proteins in oocyte competence and early development. However, their broader roles in gametogenesis, particularly in the male germline and in the regulation of germ cell homeostasis, remain poorly defined.

The *Cth1* gene encodes a CCCH zinc-finger RNA-binding protein with functional similarity to vertebrate ZFP36-family members. Cth1 and its paralog Cth2 promote post-transcriptional remodeling of gene expression during iron limitation by targeting transcripts involved in iron-consuming and mitochondrial pathways for decay or translational repression (Barlit et al, 2024; Jorda et al, 2023; Perea-Garcia et al, 2020; Ramos-Alonso et al, 2018; Romero et al, 2018). These studies suggest that Cth1-like RNA-decay factors can couple transcript turnover to metabolic adaptation and cellular homeostasis (Barlit et al, 2024; Jorda et al, 2023). Whether such mechanisms are conserved in vertebrate germ cells and contribute to fertility has remained unclear.

In zebrafish, our recent work has identified Cth1 as the first vertebrate maternally deposited RNA-decay factor whose mRNA undergoes rapid, spatiotemporally regulated decay outside the germline (Kushawah et al, 2026). We used CRISPR-Cas13d system to induce specific and robust knockdown of *cth1* transcripts in early development (Hernandez-Huertas et al, 2022; Hernandez-Huertas et al, 2025; Kushawah et al, 2020; Moreno-Sanchez et al, 2025). This promotes deadenylation-mediated turnover of maternal transcripts containing AU-rich 3′UTRs during early embryogenesis (Kushawah et al, 2026). The spatiotemporal dynamics of Cth1 are regulated by evolutionarily conserved *cis* elements within its 3′UTR. This work provided the first evidence that 3′UTR-mediated RNA localization extends beyond the well-established context of germ plasm regulation, suggesting a broader and underexplored layer of post-transcriptional control operating during early development. Although that work established a major maternal function for Cth1 during early development, adult Cth1 mutants also failed to produce viable offspring, suggesting an additional and unexplored requirement during gametogenesis. Because oogenesis, spermatogenesis, and early embryogenesis all rely heavily on post-transcriptional regulation, we hypothesized that Cth1 also acts as a germline RNA-regulatory factor required for gamete development and fertility.

Here, we investigate the gametogenic role of Cth1 in zebrafish and show that it is highly expressed in germ cells and early embryos, dynamically regulated during oocyte growth and maturation, and re-expressed upon sexual maturation. During oogenesis, *cth1* undergoes dynamic polyadenylation and spatial regulation, consistent with a role in maternal RNA control. Loss of *cth1* disrupts oocyte growth and maturation, with defects emerging during meiotic progression. We further reveal an unrecognized requirement for *cth1* during spermatogenesis. At the molecular level, Cth1 mutant germ cells exhibit transcriptomic changes consistent with metabolic and translational dysregulation, elevated DNA damage, and severe morphological abnormalities of gametes. Together, these findings support a role for Cth1 in coordinating metabolic homeostasis and genome integrity during germ cell development and fertility in both sexes.

## Results

### *Cth1* is dynamically expressed and spatially regulated during zebrafish oogenesis, spermatogenesis, and early embryogenesis

*Cth1* displays dynamic patterns of expression during the earliest stages of zebrafish development. There is a maternal deposition of transcripts in oocytes and then following fertilization, *cth1* mRNA levels rapidly decline, becoming spatially restricted to the marginal region by the mid-blastula stage. After this stage *cth1* expression is no longer detectable during early development. This dynamic expression pattern is illustrated by HCR in situ hybridization done across different developmental stages (Figures. 1A, 1B and 1C) and is further supported by analysis of publicly available bulk RNA-seq and single-cell RNA-seq datasets (Bastian et al, 2021; Sur et al, 2023). After this early embryonic phase, *cth1* expression remains largely undetectable until the onset of sexual maturation around 25 dpf (Figure. S1A). In a recent study, using SLAM-seq analysis to differentiate between maternal and zygotic transcripts (Baia Amaral et al, 2024), we demonstrated that it is the maternally deposited *cth1* transcripts that become preferentially localized to the marginal region by the mid-blastula stage (Kushawah et al, 2026; Kushawah et al, 2024). GFP: *cth1* 3′UTR reporter assays showed that reporter transcripts contain *cis*-elements that preferentially localize these transcripts to the marginal region. Hence, marginal enrichment of *cth1* is largely driven by changes in its maternally deposited mRNA and not by zygotic expression (Kushawah et al, 2026).

**Figure 1.**
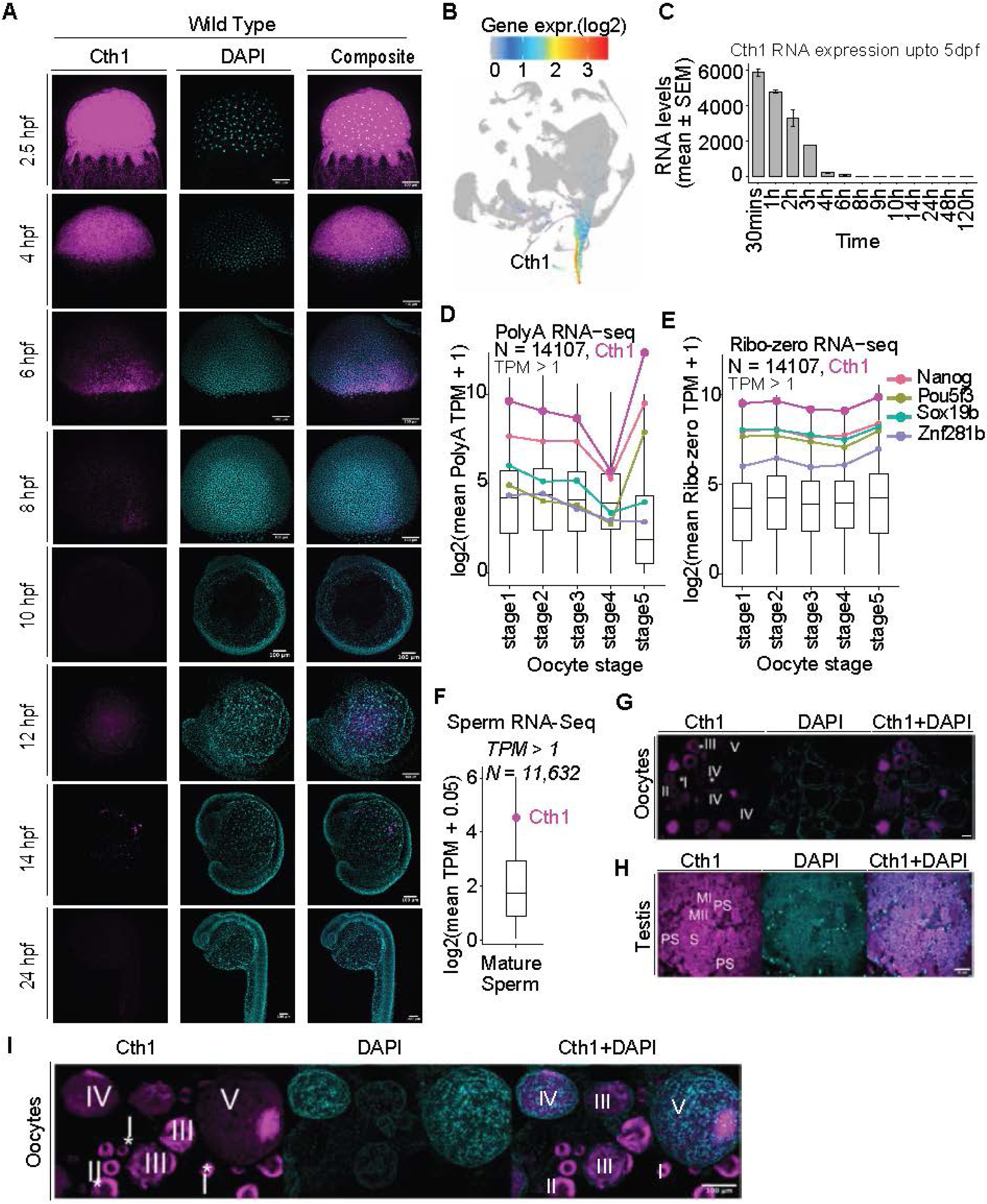
HCR and RNA-seq reveal dynamic *Cth1* RNA regulation during the gametogenesis-to-embryo transition. A Hybridization Chain Reaction (HCR) detection of endogenous *Cth1* RNA during early zebrafish embryogenesis. *Cth1* transcripts are uniformly distributed at early stages, followed by spatiotemporal enrichment at the marginal region during gastrulation, and become barely detectable by ∼8 hpf. HCR was performed on embryos from 2.5 hpf to 24 hpf. *Cth1* RNA is shown in magenta and nuclei (DAPI) in cyan. Scale bar 100 µm. B Single-cell RNA-seq data (Sur et al, 2023) from 3 hpf to 5 dpf confirm early *Cth1* expression, predominantly in blastomere cells. C Analysis of publicly available RNA-seq datasets (Bastian et al, 2021) shows high levels of *cth1* mRNA during early embryonic development, followed by rapid transcript decay by ∼4 hpf, with little to no detectable expression through 5 days post-fertilization. D Poly(A)+ RNA-seq analysis showing log mean TPM values (TPMs >1) for 14,107 genes across oocyte stages. The magenta line denotes cth1, which undergoes strong deadenylation by stage IV followed by re-adenylation at stage V. Its re-adenylation exceeds that of known zygotic genome activators (ZGA), suggesting a regulatory role during oocyte maturation and preparation for fertilization. E Ribo-depleted RNA-seq analysis showing log mean TPM values (TPMs >1) for 14,107 genes across oocyte stages. The magenta line denotes *cth1* Ribo-depleted RNA-seq of mature zebrafish sperm confirms consistent higher *cth1* expression than known zygotic genome activators. F Poly(A)+ RNA-seq analysis showing log mean TPM values (TPMs >1) for 18,300 genes in mature sperm. The magenta dot denotes *cth1*, which has higher expression levels than other expressed zygotic genome activators. G,H Representative HCR images in gonadal tissue sections of zebrafish showing robust *Cth1* (magenta) expression across different annotated gametogenic stages. *Cth1* RNA is shown in magenta and nuclei (DAPI) in cyan. Different stages in testes sections are annotated as Primordial spermatogonia (PS), Spermatogonia (S), Meiosis I (MI) and Meiosis II (MII). Scale bar 10 µm testes. I HCR analysis of zebrafish ovary showing *Cth1* (magenta) expression across oocyte stages. *Cth1* mRNA is uniformly distributed at early stages and becomes localized to the micropylar region in mature oocytes. Numbers indicate different stages of oocyte growth and maturation. Nuclei are shown in cyan (DAPI). Scale bar 100 µm.

In a previous study, we observed infertility and gametogenesis defects in *cth1* mutants (Kushawah et al, 2026) and therefore decided to examine *cth1* expression during gamete development. To assess transcriptional activity during oocyte development, we performed both poly(A)-selected and ribosomal RNA–depleted RNA sequencing across multiple developmental stages. Analysis of intronic relative to exonic reads, serves as a measure of nascent pre-mRNA abundance, and we found a marked reduction in stage V oocytes, consistent with transcriptional quiescence at this stage (Figure. S1B). Despite this global decline in transcriptional activity, *cth1* transcripts remained highly abundant throughout oocyte development (Figures. 1D and 1E). Interestingly, *cth1* transcripts undergo pronounced deadenylation by stage IV stage when meiosis I is completed, followed by rapid re-adenylation during the stage IV- V transition. By stage V, as oocytes enters meiosis II, transcripts are highly polyadenylated and remain so after arrest at metaphase II until fertilization (Figures. 1D, S1C). *Cth1* mRNA levels and polyadenylation status by stage V are comparatively higher than those for known zygotic genome activators (ZGA) and regulators, such as *nanog, pou5f3, sox19b, and* other genes (e.g. *znf281b*) which are highly deposited at 2 hpf during early development in zebrafish (Figures. 1D,1E, S1B) (Chan et al, 2019; da Silva Pescador et al, 2024; Lee et al, 2013; Medina-Munoz et al, 2021). Analyses of transcripts which similarly re-adenylated during this transition revealed enrichment for genes associated with mRNA metabolic process, DNA damage response, mRNA processing and splicing functions (Figure. S1D). Together these results suggest that Cth1 may be part of a cohort of functionally important mRNAs that are transcriptionally silent at the stage IV-V transition, when most of the maternal mRNAs undergo deadenylation (Figure. 1D).

In addition to expression in oocytes, we performed RNA seq on mature sperm collected by squeezing male zebrafish testes. This transcriptomic analysis of mature sperm further confirms that *cth1* is highly expressed in mature spermatozoa (Figure. 1F). HCR on wildtype oocytes and testes further validated *cth1* expression in both male and female zebrafish gonads (Figures. 1G, 1H, 1I, S1E, S1F, S1G,). During oogenesis, *cth1* displayed dynamic localization, showing uniform distribution at early stages and a more restricted localization in mature oocytes (Figures. 1I, S1D while in testes there was no apparent dynamics in patterns of spatial expression during different stages of spermatogenesis (Figures. 1I, S1F). Together, these results demonstrate that *cth1* is highly expressed in both gonads and dynamically regulated during female gamete development.

### Female *cth1* mutants display abnormal oocyte growth and maturation

As reported in our previous study investigating early developmental roles for maternally deposited *cth1* mRNA in early embryogenesis, we found that *Cth1* mutant females are infertile and display severe gametogenesis defects (Kushawah et al, 2026). These phenotypes were consistently observed across independently generated mutant lines using three gRNAs targeting *Cth1* gene locus individually or in combination. Here, we have used the mutant lines generated by combining all the three gRNAs to examine the potential role of *cth1* in gametogenesis, focusing on the ovarian and oocyte defects of *cth1* mutant females. Consistent with the infertility phenotype (Kushawah et al, 2026), gross anatomical analysis revealed abnormal ovary size, inflammation across different tissues, and disintegrated oocytes (Figures. 2A, 2B, S2A).

**Figure 2.**
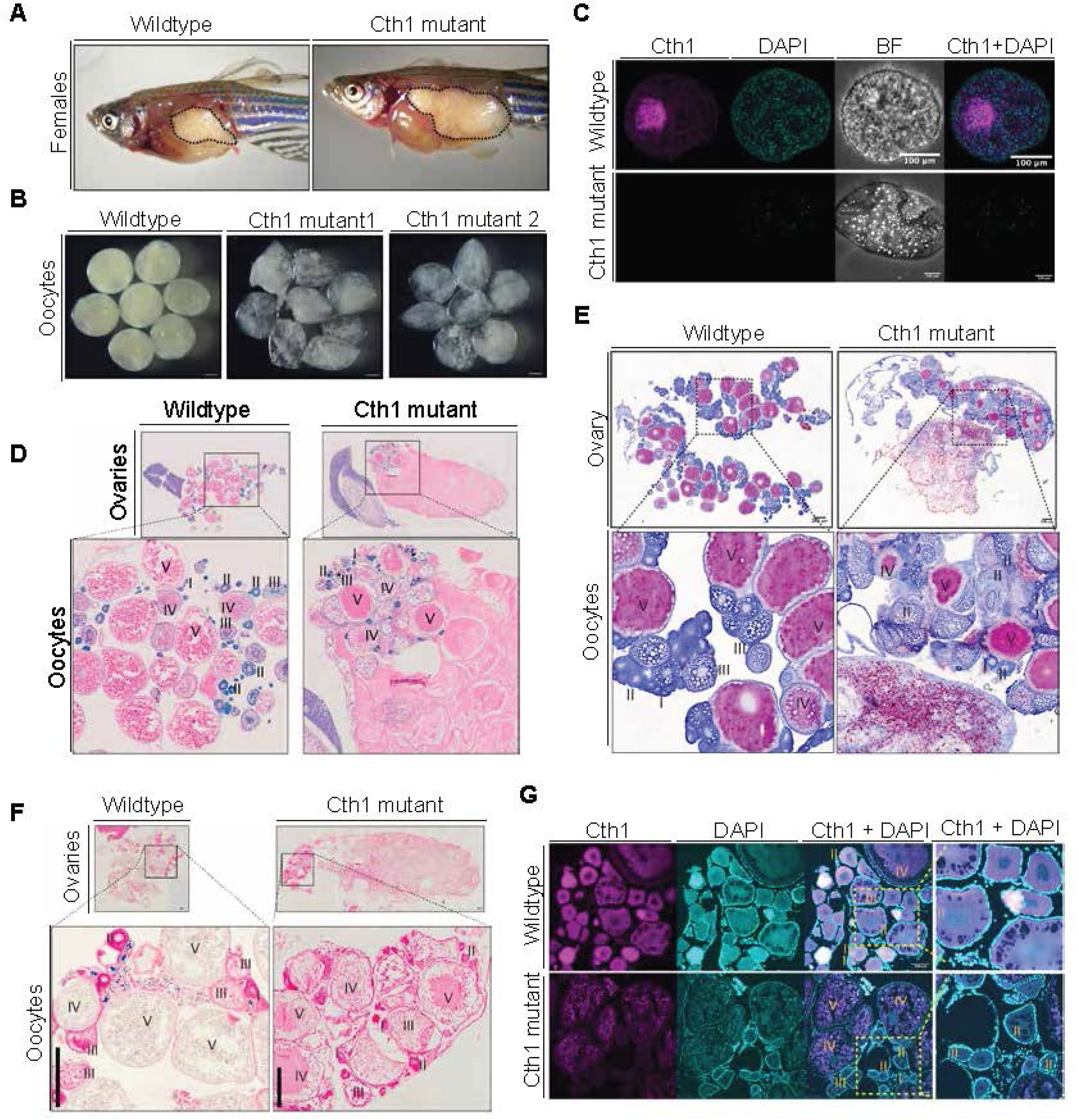
Female *cth1* mutants exhibit defective oocyte growth and maturation. A. Lateral views of wildtype and Cth1 mutant females showing internal organs. Cth1 mutant females exhibit enlarged, inflamed, and fluid-filled ovaries compared to wild type. B. Representative images of mature oocytes from Cth1 mutants appear white, translucent, and disintegrated, in contrast to intact wildtype mature oocytes. Scale bar, 100 µm. C. HCR detection of *cth1* mRNA in mature oocytes. No *cth1* transcripts signal (magenta) is detected in mature mutant oocytes, whereas wildtype mature oocytes show localized *cth1* expression. Nuclei are shown in cyan (DAPI). Low DAPI signal in mutant oocytes as they are disintegrated. Scale bar, 100 µm. D. H&E-stained ovary sections reveal the presence of dead and disintegrating oocytes in Cth1 mutant ovaries compared to wildtype. Only small regions of mutant ovaries contain growing oocytes; however, early stage I and II oocytes exhibit abnormal morphology, indicating defects in early oocyte growth. E. Oil Red O staining of freshly sectioned ovaries shows increased lipid accumulation in Cth1 mutant oocytes. By stage II, mutant oocytes contain highly stained red vesicles, suggesting altered lipid and glyceride metabolism compared to controls. Different oocytes stages are annotated. Scale bar 200 µm. F. Prussian blue staining for ferric ions (iron) reveals blue-stained particles in early-stage oocytes (cysts and stage I) in wildtype ovaries, indicating iron availability to utilize during early oocyte growth. In contrast, *cth1* mutant ovaries lack detectable Prussian blue staining at early stages and show apparently scattered blue dots in mature oocytes, suggesting impaired iron sequestration during early oogenesis. G. HCR analysis of ovary tissue sections shows downregulation of *cth1* mRNA in mutant oocytes compared to wild type. Early-stage mutant oocytes display irregular nuclear morphology and altered chromatin organization, which progressively collapse and degrade at later stages (Magnified images). Scale bar 100 µm.

Because *Cth1* is highly expressed and exhibits dynamic spatiotemporal patterning during oocyte development, we focused on the molecular and cytological consequences of *cth1* loss at successive oocyte stages. HCR analysis on mature mutant oocytes revealed a strong reduction of *cth1* transcripts, as compared to wildtype mature oocytes.

The mature mutant oocytes are highly fragmented or disintegrated and as a consequence we observed reduced DAPI signal as compared to control oocytes (Figure. 2C). Histological analysis using H&E staining revealed that mutant ovaries were largely filled with dead and disintegrated oocytes (Figures. 2D, S2B). Notably, defects were evident even at early stages of oogenesis, as stage I and stage II mutant oocytes displayed irregular morphology and intense eosin staining compared with wildtype, suggesting that Cth1 appears to be important during early stages of oogenesis (Figures. 2D, S2B). Further evidence for these early-stage abnormalities was provided by trichrome staining, which revealed excessive accumulation of cortical alveoli (dark blue vesicles) and dark cytoplasmic staining in early-stage II mutant oocytes (Figure. S2C). This reveals a disrupted cytoplasmic organization, altered cortical alveoli distribution and abnormal oocyte maturation.

To further examine these cytoplasmic changes, Oil Red O staining was performed to visualize neutral lipid accumulation. Early-stage *cth1* mutant oocytes exhibited abundant lipid droplets within the cytoplasm and intercellular spaces, consistent with altered lipid handling compared to wildtype oocytes (Figures. 2E, S2D). Prussian blue staining revealed ferric iron accumulation predominantly in the peri-oocytic/oocyte cyst region surrounding early oocytes in wildtype ovaries. In mutant ovaries, this ferric iron signal was markedly reduced/nearly absent in peri-oocytic/oocyte cyst region or showed random scattering in differentiated oocytes. These findings indicate that the mutants exhibit altered local ferric iron distribution around early oocytes, suggesting impaired iron transport within the early ovarian microenvironment (Figures. 2F, S2E). HCR analysis on sections of wildtype and mutant ovary samples containing different oocyte stages revealed early downregulation of *cth1* mRNA accompanied by abnormal chromatin organization within the nucleus at the earliest stages of oogenesis during meiosis I (Figure. 2G). As development progressed, chromatin integrity was further compromised, ultimately displaying severe chromatin disorganization and nuclear collapse in later-stage mutant oocytes compared with wildtype controls (Figure. 2G). Together, these findings indicate that loss of *cth1* disrupts oocyte development at very early stages of meiosis I, including the oocyte differentiation phase, and is likely associated with defects in cytoplasmic organization, intracellular trafficking, metabolic regulation, and nuclear integrity.

To exclude the possibility that the observed phenotypes arise indirectly from impaired homeostasis in other tissues, we generated germ cell–specific mutants using Cas9 mRNA fused to a germline-enriched *nanos* 3′UTR, which preferentially targets germ cells (Figure. S2F). Adult mutant females displayed infertility phenotypes similar to those observed in whole-body mutants (Figures. S2G, S2H), consistent with a germline-autonomous requirement for Cth1 rather than secondary effects from other tissues.

### Transcriptomic profiling of cth1 mutant oocytes reveals altered immune, translation, metabolic, and DNA-repair pathways

To define the molecular consequences of Cth1 loss, we performed RNA-seq on wildtype and Cth1 mutant mature oocytes, collected from two independent mutant lines of adult female zebrafish. As expected, *cth1* transcripts were markedly reduced in mutant oocytes. Loss of *cth1* resulted in extensive transcriptional changes, with many genes significantly upregulated in mutant oocytes (Figures. 3A, S3A).

**Figure 3.**
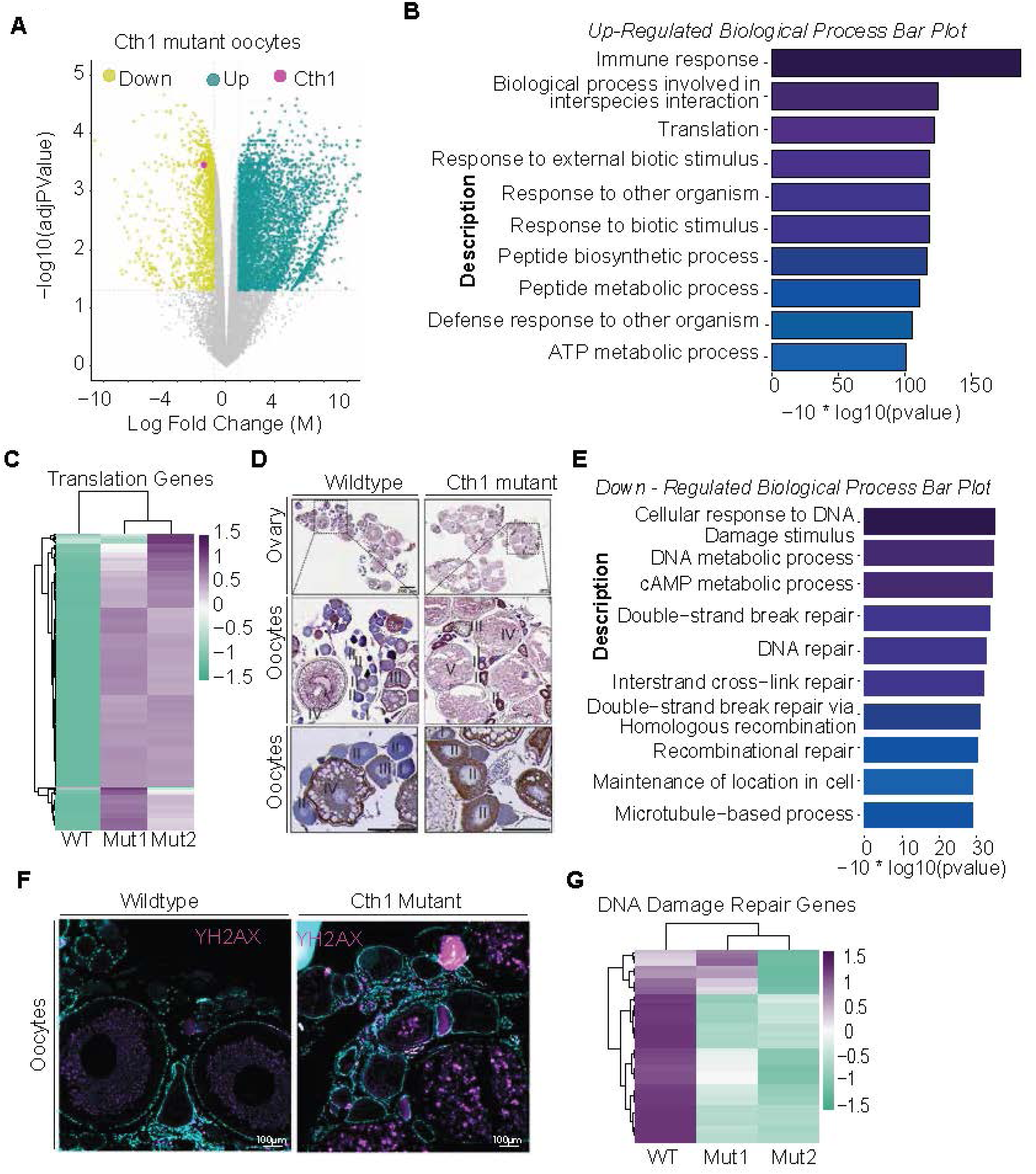
Loss of *cth1* disrupts metabolic, translational, and genome integrity pathways in oocytes. H. Volcano plot of RNA-seq analysis from mature Cth1 mutant oocytes showing adjusted *P* values and log2 fold changes relative to control oocytes. *cth1* is significantly downregulated and highlighted in magenta; differentially upregulated genes are shown in cyan and downregulated genes in yellow. I. Gene Ontology (GO) biological process enrichment analysis of upregulated genes reveals strong activation of immune response pathways, translation-related processes, and multiple metabolic pathways, including energy metabolism. J. Heatmap showing upregulation of translation-related genes from two independent *cth1* mutant oocyte samples compared to controls. K. Mitochondrial (TOM20) immunostaining of ovary sections reveals increased mitochondrial density in early-stage mutant oocytes (stage I and stage II), indicated by darker brown staining, suggesting altered metabolic activity during early oocyte growth. Scale bar 200 µm. L. GO biological process enrichment analysis of downregulated genes shows significant suppression of DNA damage repair pathways, responses to DNA damage, and cAMP metabolic processes. M. Immunostaining for the DNA damage marker γH2AX reveals elevated DNA damage in *cth1* mutant oocytes beginning at early stages of oocyte growth. Scale bar 100 µm. N. Heatmap showing downregulation of DNA damage repair genes from two independent *cth1* mutant oocyte samples compared to controls.

As described in our recent manuscript, Cth1 functions as an RNA-decay factor that targets transcripts with AU rich 3′UTRs in early developmental stages. Hence, we next asked whether transcripts upregulated in *cth1* mutant oocytes were enriched for similar motifs, as observed in early embryogenesis. However, in contrast to our previous findings from *cth1* knockdown in embryos, we did not observe enrichment of the canonical Cth1-associated 8-mer or 7-mer motifs containing AU rich sequence features (Figures. S3B, S3C). This absence of an AU motif signature may reflect that Cth1 targets a different subset of genes in oocytes versus embryos or that secondary transcriptomic changes caused by the severe developmental abnormalities observed in mature *cth1* mutant oocytes may dilute effects on primary targets.

To explore the biological processes affected by loss of Cth1, we performed gene ontology analysis on differentially expressed genes in mutant oocytes. Upregulated genes were enriched for pathways involved in immune response, translation, interspecies interaction processes, and multiple metabolic pathways, including ATP metabolism (Figures. 3B, 3C). Several of these categories, particularly immune and defense responses, may reflect the degenerating state of mutant oocytes rather than direct Cth1 regulation. These pathways were predominantly associated with cytosolic ribosomes and mitochondrial protein complexes (Figure. S3D), consistent with the increased mitochondrial abundance observed in *cth1* mutant oocytes as shown by increased levels of TOM20, a known mitochondrial marker (Figures. 3D, S3E).

In contrast, downregulated genes were highly enriched for functions related to DNA double-strand break repair and homologous recombination (Figures. 3E, 3G), as well as processes associated with nuclear chromosomes, centrosomes, and centrioles (Figure. S3F). Consistent with these findings, *cth1* mutant oocytes exhibited elevated γH2AX staining, indicating increased DNA damage (Figure. 3F). These results indicate that loss of Cth1 is associated with disrupted genome integrity, cell division, and metabolic homeostasis in oocytes.

### Loss of *cth1* disrupts spermatogenesis and male fertility

*cth1* is highly expressed during spermatogenesis (Figure. 1F) and *cth1* mutant males are infertile and exhibit severe defects in gametogenesis (Kushawah et al, 2026). Here, we examined testicular morphology, sperm production, sperm behavior, and spermatogenic defects in both type and *cth1* mutant males generated using multiple gRNAs targeting *cth1* genomic locus.

Gross anatomical analysis showed that mutant male testes were less turbid in color and displayed irregular, shrunken lobules compared with controls (Figures. 4A, 4B). Upon gentle squeezing, testes from mutant males released a watery fluid, in contrast to the turbid, milky suspension characteristic of wildtype testes, suggesting reduced sperm content in mutant male testes (Figure. S4A). Histological analysis using DAPI and H&E staining, together with electron microscopy, confirmed an obvious reduction of mature sperm within the testicular lumens of mutant male testes (Figures. 4C, 4D, S4B). Microscopic examination revealed pronounced morphological abnormalities in mutant male sperm, including fragmented or multiple flagella, rough, disintegrated head, variable thickness, and a marked reduction in sperm number (Figures. S4C, S4D). Ultrastructural analysis by electron microscopy further revealed severe defects in mutant sperm morphology, including membrane sheath disruption and abnormal flagellar structures, in contrast to the organized morphology observed in control sperm (Figure. 4E).

**Figure 4.**
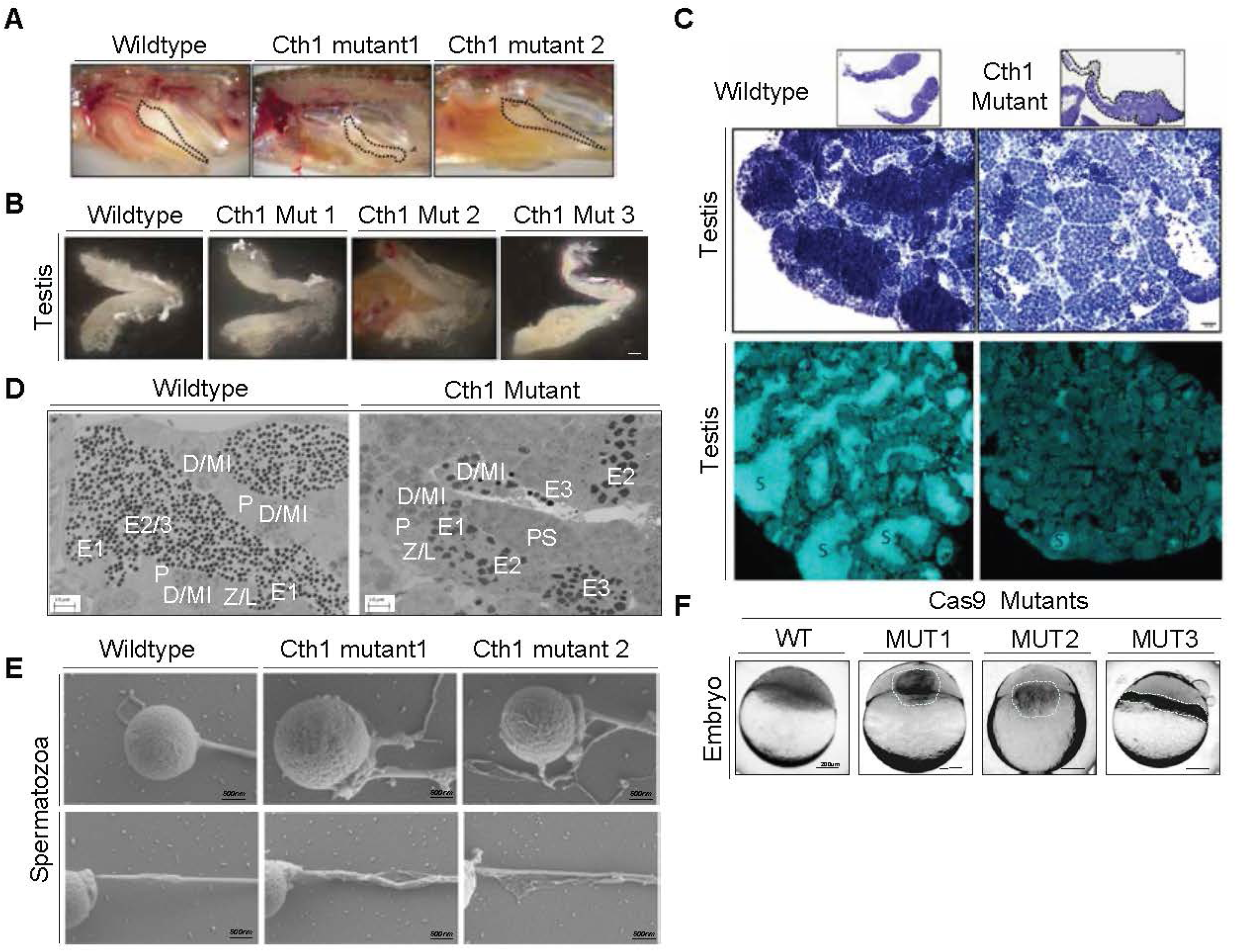
Male *cth1* mutants exhibit defective sperm growth and maturation. A. Gross morphology of testes within wildtype and *cth1* mutant males. *cth1* mutant testes appear less turbid and irregularly shaped compared to controls. B. Dissected testes from wildtype and Cth1 mutant males, wild type testes appears to have more regular lobes and more whitish in nature as compared to mutant testes from three independent mutant males. C. DAPI staining of testes sections reveals a reduced number of mature sperm in *cth1* mutant testes lobes relative to wild type. Scale bar 20 µm. D. Representative images of Scanning transmission electron microscopy of testes sections shows different stages of sperm maturation in testes lobes. Cth1 Mutant testes shows early defects during meiosis I, leading to abnormal or disintegrating sperm as compared to wildtype. Different stages of sperm growth are annotated as leptotene/zygotene (L/Z), pachytene (P), diplotene spermatocytes/metaphase I (D/MI), secondary spermatocytes/metaphase II (S/MII), early(E1), intermediate(E2), final spermatids(E3); spermatozoa (SZ). Scale bar 10 µm. E. Scanning electron microscopy of individual sperm reveals abnormal morphology in *cth1* mutant males, including multiple projections resulting in multiple tail-like structures, as well as irregular sperm heads and fragmented tail projections. Scale bar, 500 nm. F. Representative images of embryos at 4 hpf, fertilized from wildtype and Cth1 mutant male sperm. Embryos failed to fertilize using mutant male sperm as compared to wildtype male. Scale bar 200 µm.

To determine whether the fertilization defect reflects a germ cell–autonomous requirement for Cth1, rather than an indirect consequence of impaired somatic tissue homeostasis, we generated germ cell–specific Cth1 mutants by injecting multiple gRNAs targeting the *cth1* genomic locus together with Cas9 mRNA fused to the germline-enriched nanos 3′UTR (Figure. S2F). Mature sperm isolated from both whole-body and germ cell–specific cth1 mutant males fertilized eggs from wildtype adult females at severely reduced efficiency, often failing entirely. These results indicate that Cth1 is required in the male germline for fertilization (Figures. 4F, S4E).

### Transcriptomic profiling of Cth1 mutant sperm reveals altered microtubule-based processes, organelle assembly, and energy metabolism pathways

To define the molecular consequences of Cth1 loss during spermatogenesis, we performed RNA-seq on mature sperm isolated from wildtype and Cth1 mutant adult males. As expected, *cth1* transcripts were strongly reduced in mutant sperm (Figure. 5A). Loss of Cth1 resulted in widespread transcriptional changes, with a large number of genes differentially expressed (Figure. 5A). Similarly to our analyses of *cth1* mutant oocytes, mature sperm from *cth1* mutants also did not show enrichment of canonical cth1-associated 8-mer or 7-mer motifs containing AU rich sequences (Figures. S5A, S5B). Again, the lack of motif enrichment may reflect a distinct set of direct Cth1-associated targets in sperm or a broader and cumulative transcriptional perturbation in mature sperm, which could mask target-specific sequence signatures.

**Figure 5.**
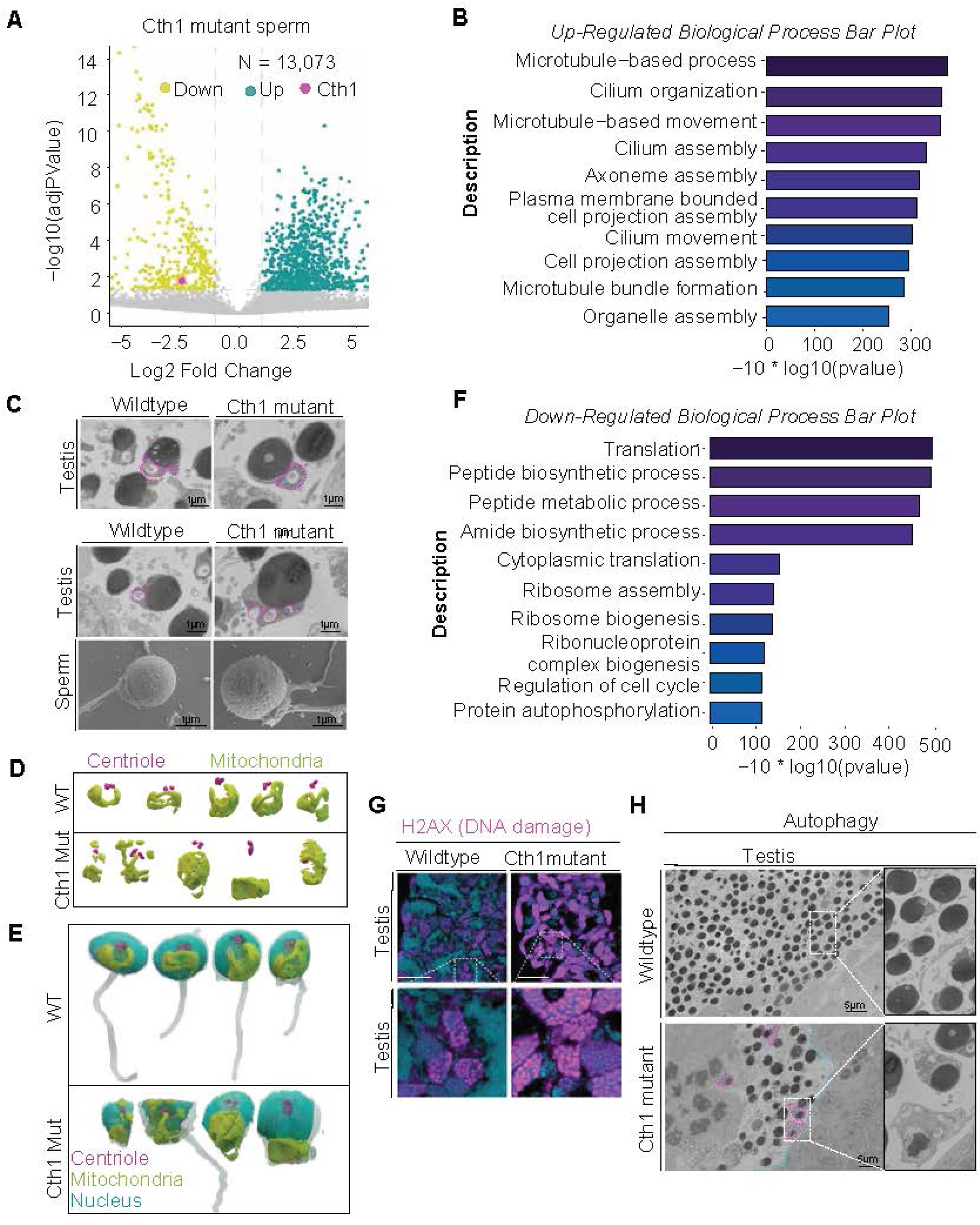
Loss of *cth1* alters transcriptional programs, genome integrity, and cellular homeostasis in mature sperm. G. Volcano plot of RNA-seq analysis from Cth1 mutant male sperm showing adjusted *P* values and log2 fold changes relative to control oocytes. *cth1* is significantly downregulated and highlighted in magenta; differentially upregulated genes are shown in cyan and downregulated genes in yellow. H. Gene Ontology (GO) biological process enrichment analysis of genes upregulated in *cth1* mutant sperm reveals enrichment for microtubule-based processes, cilium-and organelle-related organization, assembly, and motility pathways. I. Scanning and Scanning Transmission electron microscopy of testes and mature sperm reveal high density of mitochondria (Pink outline) and the presence of multiple abnormal tail projections (Pink outline) in *cth1* mutant sperm. Scale bar 1 µm. J. Three-dimensional rendering of centrioles and mitochondria segmented from serial scanning transmission electron microscopy testes sections of wildtype and Cth1 mutant males. Cth1 mutant testes show altered centriole and mitochondrial abundance and organization compared with wildtype testes. K. Three-dimensional rendering of centrioles, mitochondria, nucleus and flagella from serial scanning transmission electron microscopy testes sections of wildtype and Cth1 mutant males. Cth1 mutant testes show altered arrangement and organization compared with wildtype testes. L. GO biological process enrichment analysis of downregulated genes in *cth1* mutant sperm shows suppression of translation-related pathways, including ribosome assembly, biosynthetic processes, and ribonucleoprotein complex biogenesis. M. Immunostaining for the DNA damage markers γH2AX reveals elevated levels of DNA damage in *cth1* mutant testes compared to wildtype testes. Scale bar 100 µm. N. Scanning Transmission electron microscopy further reveals autophagosome-like structures in *cth1* mutant testes (Pink outline), accompanied by a marked reduction in mature sperm abundance compared to controls. Scale bar 5 µm.

To examine the biological processes affected by loss of Cth1, we performed gene ontology analysis on differentially expressed genes in mutant sperm. Upregulated genes were enriched for microtubule-based processes, including cilium organization, assembly, and motility, as well as organelle assembly pathways (Figures. 5B, S5C, S5D). These processes were predominantly associated with the ciliary and microtubule cytoskeleton and are consistent with the severe flagellar abnormalities observed in *cth1* mutant sperm, including multiple and disorganized tail projections (Figure. 5C). High density of mitochondrial accumulation within mutant sperm heads further supports defects in organelle organization (Figures. 5C, 5D, 5E, S5D, S5E). Together with the observed centriole alignment defects in mature *cth1* mutant sperm (Figures. 5D, 5E) these findings suggest that loss of *cth1* leads to abnormal spermiogenesis in which haploid round spermatids differentiate into mature spermatozoa. This disruption may affect centriole positioning/orientation, mitochondrial organization, and flagellum formation during sperm maturation, thereby contributing to impaired sperm development (Figures. 5D, 5E).

Downregulated genes, in contrast, were highly enriched for pathways related to translation, ribosome biogenesis and assembly, cell cycle regulation, protein autophosphorylation, and the electron transport chain (Figures. 5F, S5F, S5G). Consistent with a stressed and defective spermatogenic program, *cth1* mutant testes exhibited elevated γH2AX and RAD51 staining, indicating increased DNA damage and activation of DNA-repair pathways (Figures. 5G, S5H). Prussian blue staining revealed markedly increased ferric iron deposition in *cth1* mutant testes compared with controls, indicating altered iron homeostasis (Figure. S5I). LC3B immunostaining showed elevated autophagic activity in mutant testes, supporting increased cellular stress and turnover of defective spermatogenic cells (Figures. 5H, S5J). Ultrastructural analysis by electron microscopy further revealed the presence of potentially phagocytic structures in mutant testes, supporting enhanced autophagic like activity and consistent with increased clearance of defective germ cells (Figures. 5H, S5J). Together, these findings indicate that loss of *cth1* in the male germline is associated with broad suppression of translational and cell cycle programs, accompanied by increased genome instability and activation of germ cell clearance pathways. These defects are consistent with impaired spermatogenesis and likely contribute to the severe sperm abnormalities and male infertility observed in *cth1* mutants.

## Discussion

Gametogenesis requires the precise coordination between cellular growth, metabolic activity, cytoskeletal remodeling, and genome maintenance to generate developmentally competent gametes. Although post-transcriptional regulation is known to play a central role in germ cell development, how RNA decay pathways coordinate these processes remain poorly understood. We previously demonstrated that Cth1 functions as a spatio-temporally regulated maternal RNA decay factor promoting deadenylation-mediated turnover of transcripts which contains AU-rich 3′UTRs during early embryogenesis (Kushawah et al, 2026). Here, we extend these findings to the germline and identify Cth1 as an essential regulator of fertility in zebrafish. *Cth1* is highly enriched in oocytes, sperm, and early embryos and undergoes dynamic spatial, temporal, and poly(A)-tail regulation during oogenesis and early embryogenesis (Figure. 1). In contrast, although *cth1* is abundant in mature sperm, its spatiotemporal and poly(A)-tail regulation during spermatogenesis will require further investigation. Further, *cth1* becomes one of the most highly adenylated transcripts in mature, transcriptionally silent oocytes, showing higher adenylation than known zygotic genome activators (Figure. 1). Similarly in mature sperm *cth1* transcripts levels were high. This suggests that *cth1* may be under precise post-transcriptional control and is potentially primed for rapid translation after fertilization before the acquisition of developmental competence (Figure. 1). Consistent with these expression dynamics, loss of *cth1* causes complete infertility in both sexes and disrupts multiple aspects of germ cell biology, including metabolic homeostasis, cytoskeletal organization, and genome integrity (Figures. 2-5). Together, these findings support a model in which Cth1 contributes to the maternal RNA post-transcriptional regulatory network linking gametogenesis to early embryonic development and highlight RNA decay as an important regulator of reproductive success.

In oogenesis, our data suggest that Cth1 restrains metabolic and translational programs during early oocyte development, particularly at stages I–II, when oocytes undergo meiotic progression and extensive nuclear remodeling (Figure. 2). *Cth1* mutant oocytes display early cytoplasmic disorganization, altered intracellular trafficking, increased cortical alveoli accumulation, elevated mitochondrial content, abnormal lipid accumulation, disrupted ferric iron distribution, and progressive chromatin collapse (Figure. 2). These defects suggest that loss of Cth1 disrupts the balance between cellular growth, metabolic control, and nuclear integrity during early oogenesis. Consistent with this model, mutant oocytes show downregulation of DNA repair pathways and accumulation of DNA damage markers, indicating that genome maintenance becomes uncoupled from growth-associated programs in the absence of *cth1* (Figures. 2 and 3). Transcriptomic analyses further support this hypothesis by revealing widespread upregulation of metabolic, mitochondrial, proteolysis, and translational pathways alongside repression of cell cycle and genome maintenance programs. These opposing transcriptional trends suggest a model in which Cth1 may normally help enforce a temporal separation between growth and genome protection, a mechanism previously implicated in oocyte maturation but not mechanistically defined. Our findings extend prior work in *Xenopus* by proposing that, Cth1 may contribute to translational timing but also maintaining nuclear stability during oogenesis (Belloc & Mendez, 2008).

In spermatogenesis, loss of *Cth1* causes profound defects in testes morphology, sperm production, and fertilization capacity. *Cth1* mutant testes show impaired spermatogenesis, resulting in reduced mature sperm and abnormal sperm morphogenesis, including defective flagellar assembly and disrupted sperm head and tail architecture (Figure. 4). RNA-seq analysis of mutant sperm reveals enrichment of microtubule- and cilium-associated programs, consistent with dysregulated cytoskeletal remodeling during spermiogenesis (Figure. 5). These molecular changes align with the abnormal flagellar duplication, extension, and disorganization observed in mutant sperm. In parallel, downregulation of translational and cell cycle pathways, together with increased DNA damage, autophagy, and phagocytic activity, suggests that defective spermatogenic cells fail to complete maturation and are instead targeted by germline quality-control pathways.

Altered ferric iron distribution in mutant ovaries and testes, further suggests that Cth1 loss perturbs iron-associated metabolic homeostasis, although further detailed studies are needed to decipher the mechanism. Because iron is essential for mitochondrial function but can also promote oxidative stress when mis-regulated, disrupted iron handling may contribute to the metabolic imbalance and genome instability observed in *cth1* mutants. In Yeast, Cth1 is known to sense the iron deficiency in the growth media and help to adapt to the environment by halting the non-essential, high iron consuming processes like mitochondrial respiration, lipid biosynthesis (Barlit et al, 2024; Ramos-Alonso et al, 2018; Romero et al, 2018). Although the direct relationship between Cth1-dependent RNA regulation, iron metabolism, and DNA damage remains to be defined in zebrafish, these findings raise the possibility that impaired iron buffering amplifies mitochondrial and nuclear stress during gametogenesis.

Together, these findings support a model in which Cth1 integrates RNA decay, metabolic restraint, cytoskeletal organization, and genome protection to promote successful gametogenesis. This work positions Cth1 as a key regulator of germline quality control and suggests that disruption of RNA-based regulatory mechanisms may be an important mechanism underlying infertility. The severe phenotypes observed in *cth1* mutants during early gametogenesis highlight an essential role for Cth1-dependent RNA regulation in germline development. However, the early onset of these defects, together with the technical difficulty of isolating early-stage gametes from zebrafish at reproductive maturity, makes it challenging to define the primary molecular targets of Cth1 using conventional approaches. Future studies will therefore require carefully staged analyses at the onset of sexual maturity to identify direct Cth1-bound transcripts in both oocytes and sperm. Integrating RNA-seq with complementary RNA–protein interaction approaches will be critical to determine how Cth1-regulated RNA networks control mitochondrial activity, lipid metabolism, cytoskeletal remodeling, DNA repair, and other processes required for gamete quality.

An important future direction will also be to determine how *cth1* itself is dynamically regulated during gametogenesis and early embryogenesis. Given the established roles of 3′UTR *cis*-regulatory elements in controlling RNA stability, localization, poly(A)-tail length, and translation, reporter zebrafish lines carrying wildtype or mutant *cth1* 3′UTRs would provide a powerful *in vivo* platform. These lines would enable tissue-specific tracking of *cth1* mRNA and protein localization, decay, polyadenylation, and clearance across oocyte maturation, fertilization, and early development. Combined with CLIP-seq or RNA immunoprecipitation, poly(A)-tail profiling, and 3′UTR mutagenesis, these approaches will help distinguish direct Cth1 targets from secondary stress responses and define how Cth1-dependent RNA regulation safeguards germline development and reproductive competence in zebrafish. Together, these future studies will provide a framework for understanding how post-transcriptional regulation preserves gamete integrity and supports the successful transition to embryonic development.

## Lead Contact

Further information and resource request should be directed to lead contacts Gopal Kushawah and Ariel A. Bazzini.

## Data and Code Availability

The Accession number for the RNA-seq data used in this manuscript is GEO: XXXX. All relevant data is available upon request from the corresponding author.

## Declaration of generative AI and AI-assisted technologies in the writing process

During the preparation of this work the authors used ChatGPT and Claude in order to improve only the text. After using this tool, the authors reviewed and edited the content as needed and take full responsibility for the content of the publication.

## Acknowledgments

We thank Robb Krumlauf and Cameron Berry for critical reading and valuable suggestions on this manuscript; the core facilities at the Stowers Institute, including Sequencing and Discovery Genomics, Genome Engineering, Automation and PCR Technology, and Media Prep; and all members of the Bazzini laboratory for their intellectual and technical support. Thank you to Melainia McClain for assistance with serial section image alignment and Xia Zhao for EM screening. This study was supported by the Stowers Institute for Medical Research. AAB was awarded a Pew Innovation Fund and US National Institutes of Health grants (NIH-R01 GM136849, NIH-R21 OD034161 and NIH-1R35GM161459). This work was performed as part of post-doctoral training for GK at the Stowers Institute for Medical Research. Original data underlying this manuscript can be accessed from the Stowers Original Data Repository at http://www.stowers.org/research/publications/libpb-XXXX.

## Author contributions

G.K. and A.A.B. conceived the project and designed the research.

G.K. performed all zebrafish experiments.

G.K. performed data analysis.

S.N. performed electron microscopy.

C.C. performed sperm and different stages of oocytes collection.

M.S. and H.W. performed histology and tissue staining experiments.

N.Z. performed data analysis on different stages of oocytes.

G.K. and A.A.B. wrote the manuscript with input from the other authors. All authors reviewed and approved the manuscript.

## Methods

### Zebrafish maintenance, embryo productions and experimentations

All experiments involving zebrafish (*Danio rerio*) were conducted under protocols approved by the Institutional Animal Care and Use Committee (IACUC) at the Stowers Institute. Embryos were obtained from adult AB, TU, TF, and TLF zebrafish strains aged 6–18 months. Breeders were randomly selected from a colony of approximately 500 fish representing four independent genetic backgrounds. Embryos were cultured in standard zebrafish embryo medium at 28.5°C under routine laboratory conditions.

### Hybridization Chain Reaction (HCR) for zebrafish embryos and tissues

Custom HCR probes against *cth1*, *trim8a*, and *eno4*, together with the appropriate amplifiers and reagents, were purchased from Molecular Instruments. Whole-mount HCR and tissue specific HCRs were carried out according to the manufacturer’s protocol for zebrafish embryos. During the amplification step on day 2, fluorophore-conjugated amplifiers were added, and samples were incubated overnight at room temperature protected from light. Embryos were then washed, counterstained with DAPI, and prepared for imaging.

### Gamete collection from adult zebrafish

Gametes were obtained from 6–9-month-old wildtype and mutant adult zebrafish in accordance with institutional animal care and handling regulations. Different stages of oocytes for RNA-seq were collected following protocol (Elkouby & Mullins, 2017). To collect mature oocytes, females were placed with males overnight in breeding tanks separated by a divider. The next morning, at the onset of the light cycle, females were anesthetized in 4 g/L MS-222, and mature oocytes were collected by gentle abdominal stripping. For sperm collection, males were anesthetized under the same conditions and positioned ventral side up in a fish holder. Sperm was then collected using a 10 µL microcapillary pipette attached to an aspirator tube assembly.

### Histology, tissue staining, immunostaining, and HCR on gonadal sections

Fresh and/or fixed tissues were collected from adult wildtype and *cth1* mutant animals and processed according to the requirements of each downstream staining procedure. For histological analyses, tissues were fixed, embedded, sectioned, and subjected to standard staining protocols, including hematoxylin and eosin (H&E) staining for general tissue morphology, Masson’s trichrome staining for connective tissue and fibrosis assessment, Oil Red O staining for neutral lipid detection, and Prussian Blue staining for ferric iron accumulation. For protein detection, immunostaining was performed on fixed and processed ovary and testes sections from wildtype and cth1 mutant samples in parallel following the respective established protocols and manufacturer’s recommendations.

### RNA isolation, cDNA synthesis, and quantitative RT–PCR

To assess transcripts levels, total RNA was extracted from embryos, oocytes and mature sperm using either TRIzol reagent (Thermo Fisher Scientific) or the Zymo Research RNA Mini Kit (Zymo Research, Cat. No. R1055), according to the manufacturers’ instructions. RNA concentration was determined using the Qubit RNA Broad Range assay. For quantitative RT–PCR analysis, cDNA was generated using the SuperScript IV First-Strand Synthesis System (Thermo Fisher Scientific, Cat. No. 18091050) following the manufacturer’s protocol. Primers used for amplification are listed in Table S1. qRT–PCR reactions were prepared using low ROX SYBR Green dye (QuantaBio, Cat. No. 95074-05K), gene-specific primers at 20 μM, and 1–2 μl of diluted cDNA in a final reaction volume of 25 μl. Reactions were assembled using an automated Freedom EVO® PCR workstation (Tecan) with a predefined template program according to sample number. Target transcript levels were normalized to endogenous reference genes, including cdk2ap2 or taf15.

### Scanning Electron Microscopy

Sperm samples were diluted and applied to coverslips pre-coated with 4 nm gold–palladium using a Leica EM Ace600. Samples were prepared using a membrane-preserving protocol adapted from Korneev, Merriner et al. (2021). After critical point drying with a Tousimis Samdri-795, samples were coated with an additional 4 nm layer of gold–palladium. Imaging was performed on a Zeiss Merlin SEM using an SE2 detector at 3kV and 100 pA. Images were processed uniformly in Fiji using the CLAHE plugin (Schindelin et al, 2012).

### Scanning Transmission Electron Microscopy (STEM)

Testes samples (<1mm^3^) were fixed in 0.1M sodium cacodylate buffer with 2.5% glutaraldehyde, 2% paraformaldehyde, 1mM CaCl2 and 1% sucrose for 1 h at room temperature. The fixative was then refreshed, and samples were held overnight at 4 °C on a nutator. After four, 10-min buffer rinses, samples were post-fixed in 1% buffered osmium tetroxide for 1 h, followed by buffer rinses as before. Samples were then washed four times for 10 min in ddH₂O and incubated in 0.5% aqueous uranyl acetate overnight at 4 °C on a nutator. The next morning, after four 10-min ddH₂O washes, samples were dehydrated through a graded ethanol series (30, 50, 70, 80, 90%) followed by four 100% ethanol changes, then infiltrated with Hard Plus 812 resin (Electron Microscopy Sciences) at 25% (30 min), 50% (1 h), 75% (1 h), and 100% (2 h), followed by 100% overnight and a final 100% exchange containing accelerator the next morning. Samples were embedded in flat molds and polymerized at 60 °C for 48 h.

Because of low mature sperm counts, cth1 mutant blocks were first screened by toluidine blue staining followed by brightfield imaging to identify regions of interest (ROIs) for serial sectioning. Blocks were serially sectioned at 80 nm on a Leica UC7 ultramicrotome with a Diatome Ultra 45° diamond knife and collected on Formvar/carbon-coated slot grids (Electron Microscopy Sciences). Grids were post-stained with Sato’s lead (Sato, 1968) (3 min), 1% uranyl acetate (4 min), and Sato’s lead (5 min), then dried. Serial sections were imaged at 5 nm XY resolution on a Zeiss Merlin SEM with a STEM detector at 28 kV and 700 pA using Atlas software. Datasets were aligned in IMOD (Kremer et al., 1996) with Midas, applying local warping corrections focused on regions containing mature sperm. Mature sperm that were fully or mostly contained within the imaged volume, and in which the flagella and centrioles could be assessed, were manually segmented in IMOD (nucleus; cell body, including flagellum; and centrioles). Segmentations were meshed and rendered in Blender v4.0.(Community, 2018)(Kremer et al, 1996; Sato, 1968).

### Light Microscopy Image acquisition

Images were acquired using different imaging platforms according to sample type and experimental requirements. Overview images of zebrafish embryos at early developmental stages up to 1 dpf were captured using a Leica stereo-fluorescence microscope equipped with a DFC900 camera. Representative bright-field images of individual embryos were acquired using a Leica MZ APO stereo microscope. HCR samples were imaged on a Nikon spinning-disk confocal microscope using 10X and 20X objectives. Most of the histology tissue sections were imaged using Olympus VS120 Slider Scanning System using 5X and 10X magnification. Images of adult fish were taken using a Samsung Galaxy S21 Ultra camera. Sperm samples were imaged by scanning electron microscopy using a Zeiss Merlin SEM equipped with an SE2 detector. Image processing was performed in Fiji, and final figure panels were assembled in Adobe Illustrator 2026.

### RNA-seq libraries preparation and primary analysis

Total RNA was extracted from oocyte and sperm samples obtained from wildtype and Cth1 mutant animals. For each oocyte sample, RNA was isolated from 50 oocytes. For sperm samples, material was collected from three adult males per sample. RNA extraction from oocytes was performed using TRIzol reagent (Thermo Fisher Scientific), and from sperm RNA was isolated using PureLink^TM^ RNA mini kit (Invitrogen) and RNA quality was assessed using a Bioanalyzer (Agilent). For oocyte RNA-seq library preparation, only samples with an RNA integrity number (RIN) greater than 7.5 were used. RIN values were not used as a selection criterion for sperm RNA samples. For library preparation, at least 100 ng of total RNA from oocytes and 10–20 ng of total RNA from sperm were used with reagents compatible with the NEBNext Ultra II workflow and Nextera-type adapters. Libraries were sequenced as 100-bp single-end reads on the G4-F3 platform using a Singular Genomics flow cell (Cat. No. 700125). Sequencing reads generated on the G4-F3 platform were demultiplexed into FASTQ files using Singular Genomics sgdemux v1.2.0, allowing up to one mismatch. Reads were then aligned to the zebrafish GRCz11 reference genome from Ensembl using STAR v2.7.10b. Gene-level read counts were generated using the Ensembl gene annotation release 106. Transcript abundance was quantified as transcripts per million (TPM) using RSEM v1.3.1.

### Data Analysis

RNA-seq processing and downstream analyses were performed in R using packages from CRAN and Bioconductor. Differential expression analysis was carried out in R v4.2.3 using edgeR v3.40.2. Reads were analyzed against the zebrafish danRer11/GRCz11 reference genome with Ensembl gene annotation release 106. Gene-level count matrices were used for downstream statistical analyses. Plots were generated using ggplot2, and gene ontology analyses were performed using appropriate zebrafish-compatible Bioconductor annotation packages.

---

**Supplementary Figure S1.**
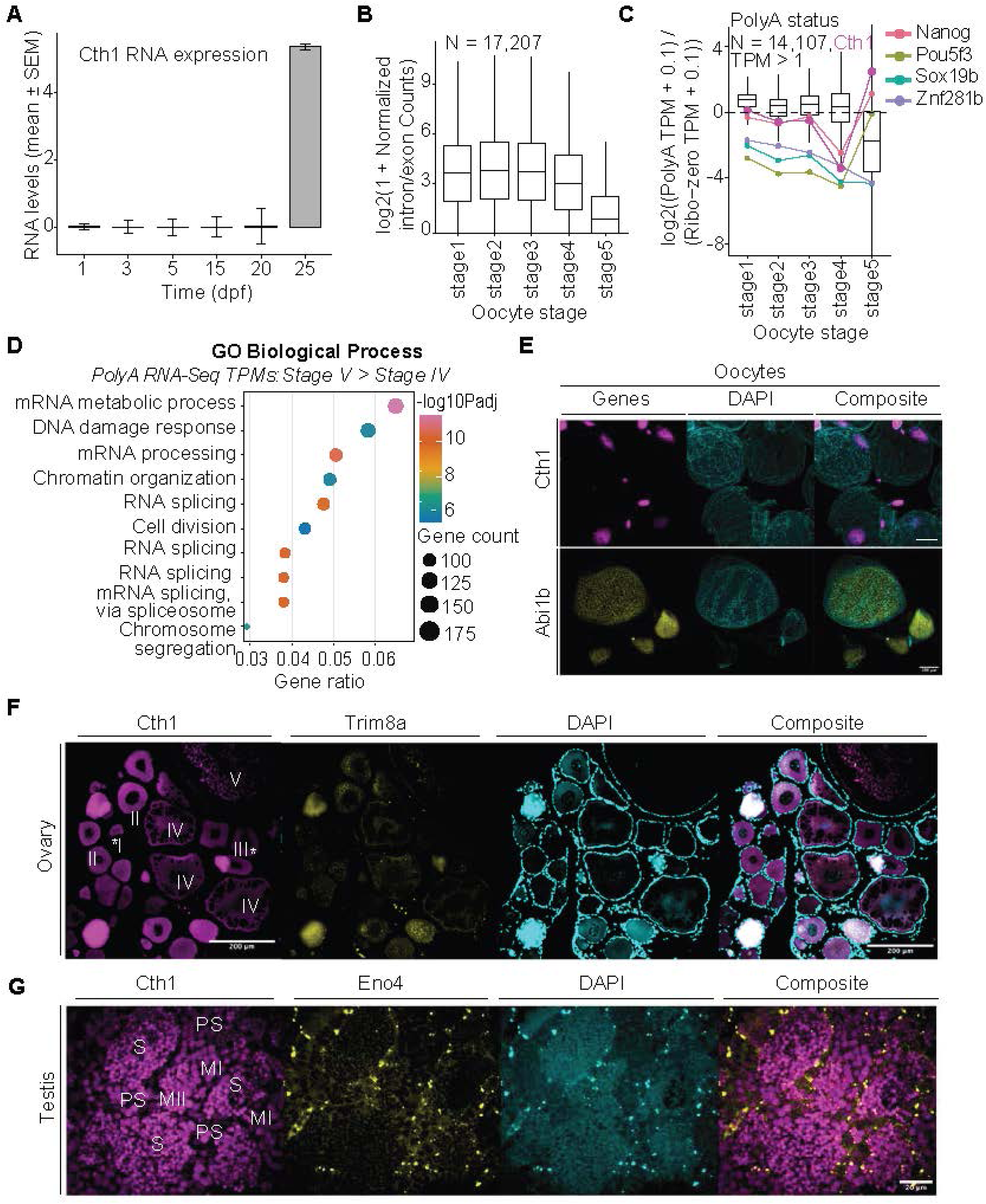
HCR and RNA-seq reveal dynamic *cth1* RNA regulation during the gametogenesis-to-embryo transition. A. The qRT-PCR analysis of *cth1* expression from 1 dpf to 25 dpf shows re-expression at 25 dpf, coinciding with sexual maturation. B. Plot showing the relative abundance of nascent transcripts across oocyte developmental stages, estimated from the ratio of intronic reads, representing pre-mRNA, to exonic reads. Stage V oocytes show a substantially lower intron-to-exon ratio compared with earlier developmental stages, indicating reduced production of newly transcribed RNA and suggest transcriptional silencing of mature, fertilization-competent oocytes stage V. C. Plot showing Poly(A) status, calculated as the log fold change between Poly(A)+ and ribo-depleted RNA-seq TPMs for 14,107 genes with TPM > 1. The magenta line denotes *cth1*, which shows low poly(A) status till stage IV followed by strong polyadenylation at stage V, reaching levels higher than known zygotic genome activators. This suggests a potential regulatory role for *cth1* during oocyte maturation and preparation for fertilization. D. Plot showing Gene Ontology (GO) pathways associated with biological process enrichment analysis of genes with TPM > =1 which gets poly adenylated from stage IV-V transition. E. HCR detection across oocyte developmental stages reveals dynamic spatial and temporal localization of *cth1*(magenta), with enrichment in mature oocytes. HCR for *abi1b* (yellow) was used as a control to demonstrate that the observed *cth1* localization is not an artifact of the assay. Scale bar 100 µm. F,G. HCR analysis of ovary and testes tissue sections confirms robust *cth1* expression in both tissues (magenta). *trim8a* (ovary) and *eno4* (testes) were used as positive controls. Nuclei are shown in cyan (DAPI). Scale bars, 200 µm (ovary) and 20 µm (testes).

**Supplementary Figure S2.**
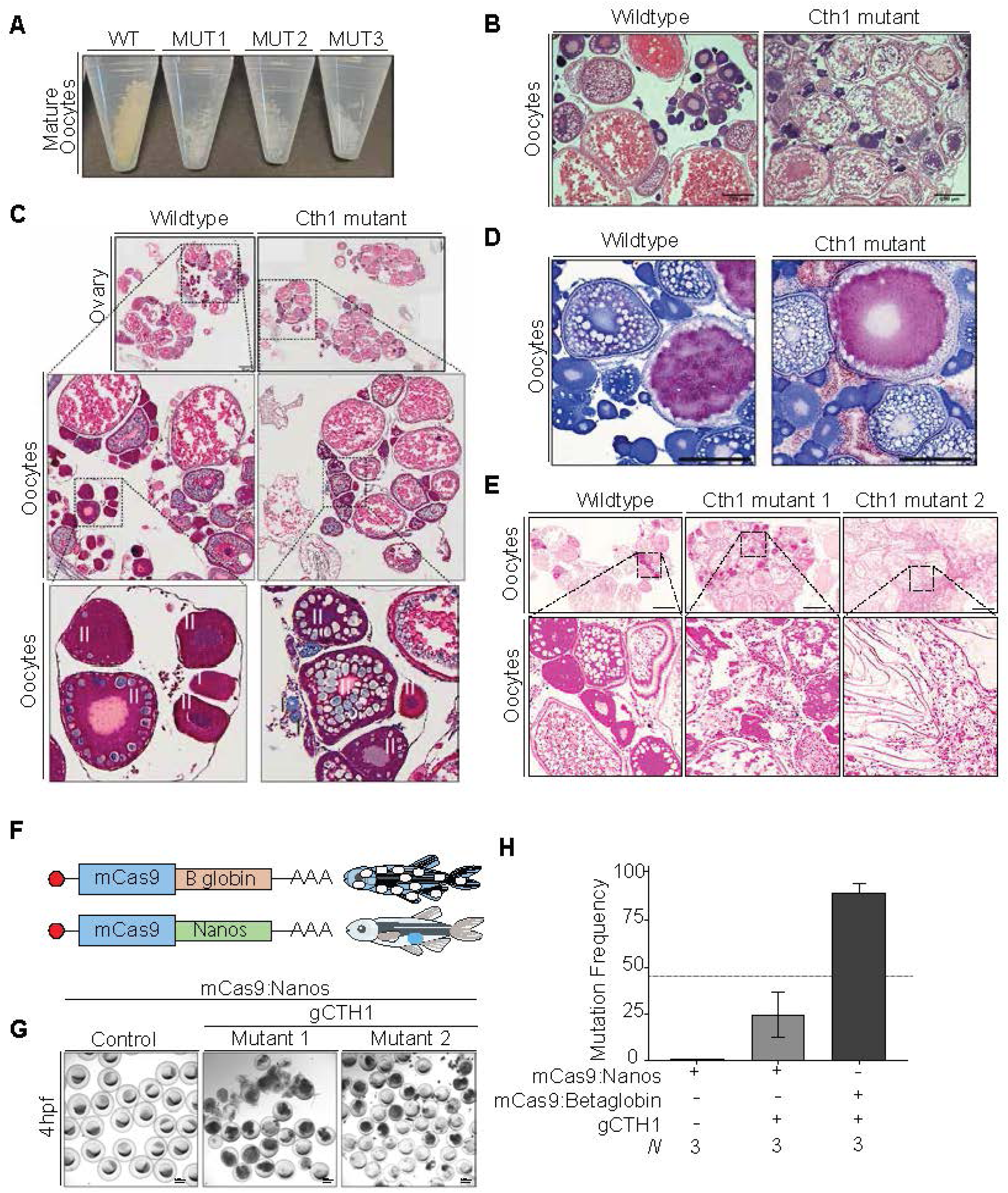
Female *cth1* mutants exhibit defective oocyte growth and early metabolic alterations. A. Mature oocytes collected by stripping from adult female zebrafish. Oocytes from *cth1* mutant females appear white and opaque, whereas oocytes from wildtype females are translucent and pale yellow. B. H&E-stained ovary sections reveal altered ovarian architecture in *cth1* mutant females, with abnormal oocyte morphology evident from early stages of oocyte growth. Many stage II oocytes display irregular nuclear morphology. Scale bar, 200 µm. C. Trichome staining of ovary sections shows increased staining intensity and enlarged germinal vesicles in early-stage *cth1* mutant oocytes compared to controls, suggesting altered metabolic or cytoplasmic organization during early oogenesis. Scale bar 200 µm. D. Oil Red O staining of freshly sectioned ovaries reveals increased accumulation of lipid and triglyceride-rich particles in stage II *cth1* mutant oocytes, indicating disrupted lipid metabolism during early oocyte development. Scale bar 200 µm. E. Prussian blue staining for ferric ions (iron) reveals blue-stained particles in early-stage oocytes (cysts and stage I) in wildtype ovaries, indicating iron availability to utilize during early oocyte growth. In contrast, *cth1* mutant ovaries lack detectable Prussian blue staining at early stages and show apparently scattered blue dots in mature oocytes, suggesting impaired iron sequestration during early oogenesis. Scale bar 100 µm. F. Cartoon showing the strategy to generate germ cell specific Cas9 mutants. G. Representative embryos from germ cell specific mutant females when crossed with wildtype male. F1 embryos produced by crossing mutant females were either unfertilized or disintegrated as compared to wildtype females. Scale bar 500 µm. H. Fin clip genotyping from germ cell specific mutants shows low mutation frequency as compared to whole body mutant. N, fin clips were used.

**Supplementary Figure S3.**
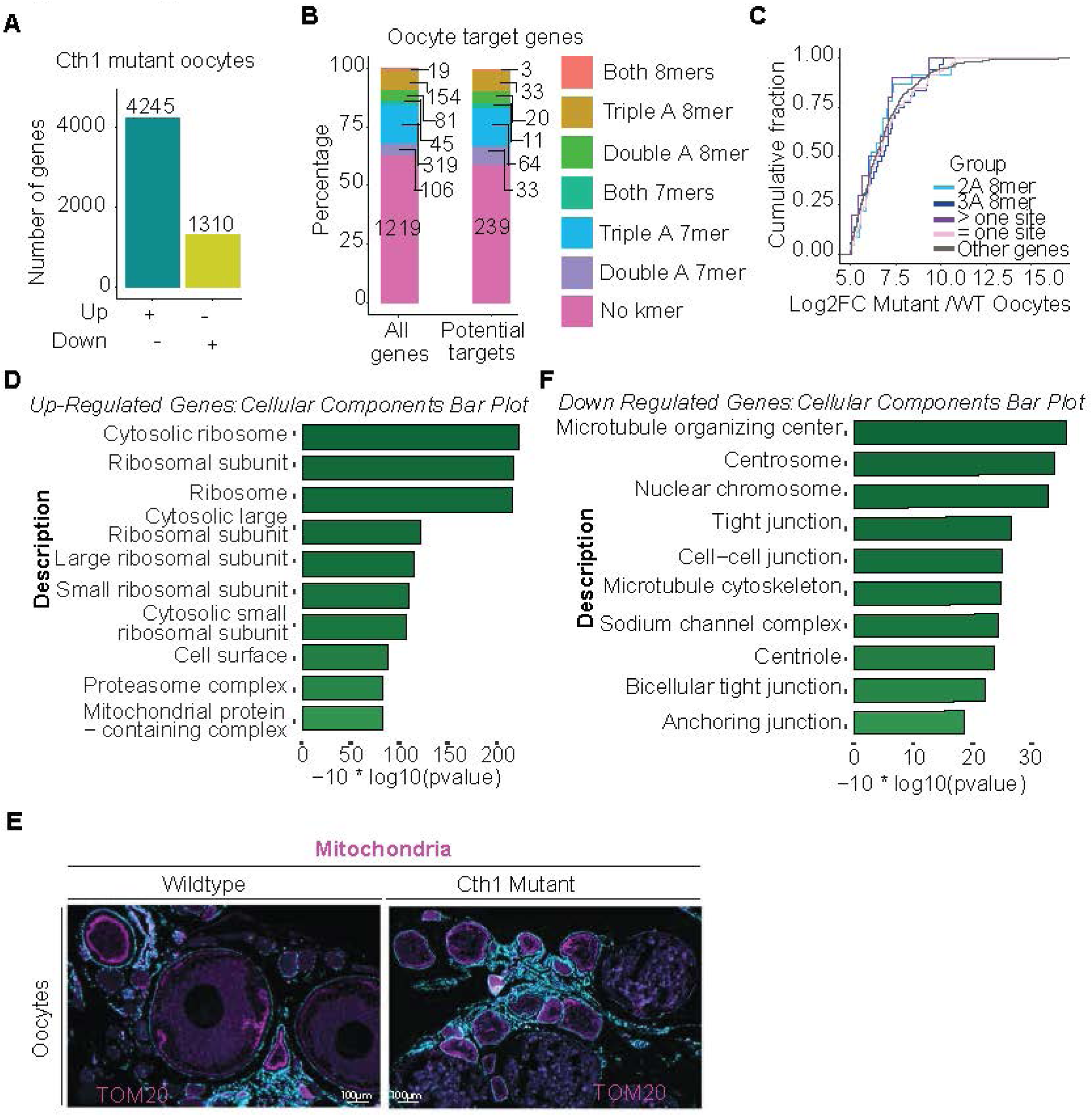
Loss of *cth1* disrupts metabolic, translational, and genome integrity pathways in oocytes. A. Bar plot showing the number of gene differentially regulated in Cth1 mutant oocytes as compared to wildtype fish. B. Percentage of the 7-mer and 8-mer motifs containing “AAA” and “AA” one or more times in the putative Cth1 target and all the genes. The putative Cth1 target group genes with enriched motifs with these combinations apparently do not show big difference as compared to all the genes constitution. C. Cumulative plot showing fold change between Cth1 mutant and wildtype oocytes. Transcripts containing enriched 7-mers/8-mers, “AAA” and multiple motifs, do not exhibit greater destabilization than other genes. D. Gene Ontology (GO) cellular component enrichment analysis of differentially expressed genes in Cth1 mutant oocytes. Upregulated genes are enriched for ribosomal, mitochondrial, and proteasome-related complexes. E. Immunostaining for the mitochondrial marker TOM20 reveals increased mitochondrial density in Cth1 mutant oocytes compared to wildtype controls. Scale bar 100 µm. F. Gene Ontology (GO) cellular component enrichment analysis of downregulated genes is associated with nuclear chromosome, centrosome, centriole, and microtubule cytoskeleton components.

**Supplementary Figure S4.**
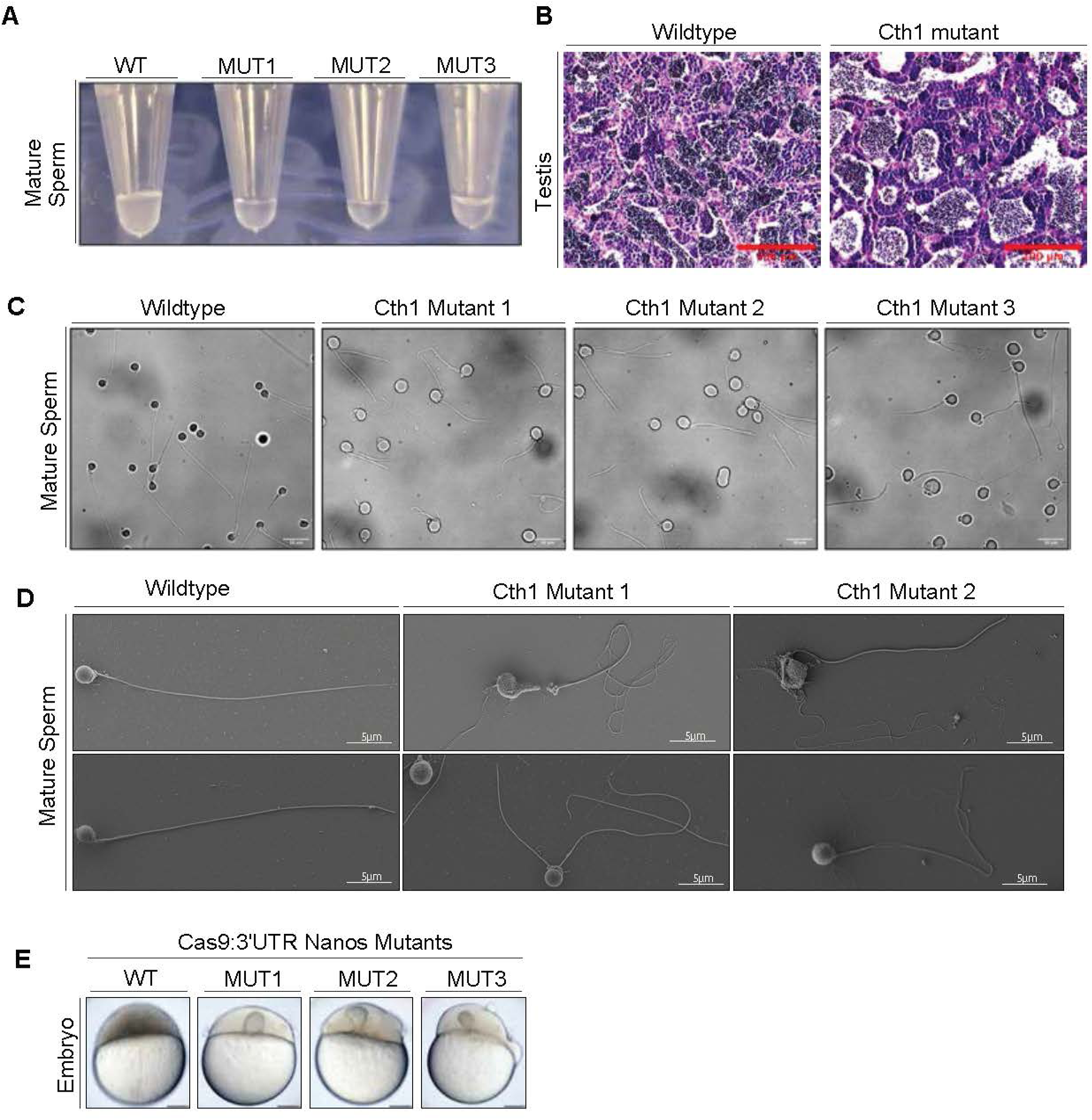
Male *cth1* mutants exhibit defective sperm development and fertilization capacity. A. Mature sperm collected by gentle abdominal pressure show reduced turbidity and a more dilute appearance in Cth1 mutant samples compared to the opaque, turbid sperm suspensions from wildtype males. B. H&E staining on representative testes sections show low number of mature sperm in Cth1 mutant as compared to wildtype testes. Scale Bar 100 µm. C. Confocal images of mature sperm at 100× magnification reveals abnormal sperm morphology in Cth1 mutants, including irregular head shape, multiple or fragmented flagella, or complete absence of flagella. Scale bar, 10 µm. D. Scanning electron microscopy of mature sperm from wildtype and independent *cth1* mutant males. Mutant sperm display pronounced structural abnormalities, including wrinkled and degraded heads and variable tail defects, ranging from absent or shortened tails to multiple fragmented tail structures. Scale bar, 5 µm. E. *In vitro* fertilization assays using mature sperm from germ cell specific Cth1 mutant males to fertilize wildtype eggs demonstrate severely reduced fertilization efficiency, resulting in either failed or incomplete fertilization compared to controls. Embryos are shown at 3 hpf. Scale bar 100 µm.

**Supplementary Figure S5.**
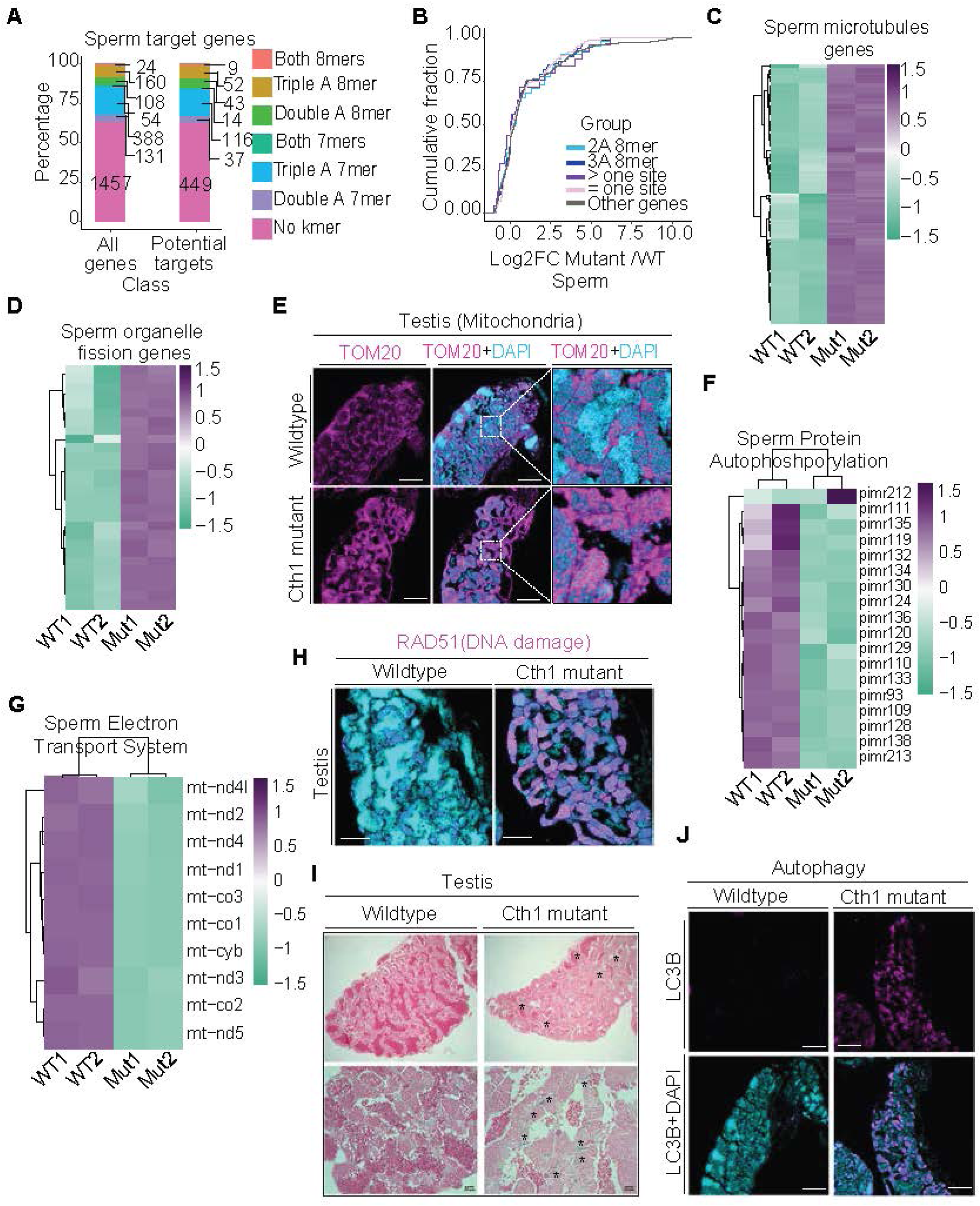
**Loss of *cth1* alters transcriptional programs, genome integrity, and cellular homeostasis in mature sperm**. A. Percentage of the 7-mer and 8-mer motifs containing “AAA” and “AA” one or more times in the putative Cth1 target and all the genes in mutant sperm. The putative Cth1 target group genes with enriched motifs with these combinations apparently do not show big difference as compared to all the genes constitution. B. Cumulative distribution of transcript fold changes between *cth1* mutant and wildtype sperm. Transcripts containing enriched 7-mer or 8-mer motifs, including “AAA” and repeated motif classes, do not exhibit greater destabilization than the remaining genes. C. Heatmaps from RNA-seq analysis of Cth1 mutant sperm show upregulation of genes associated with sperm microtubule-based processes compared to wildtype controls. D. Heatmaps from RNA-seq analysis of Cth1 mutant sperm show upregulation of genes associated with organelle fission compared to wildtype controls. E. Immunostaining for the mitochondrial marker TOM20 reveals increased mitochondrial density in Cth1 mutant testes relative to wild type. Scale bar 100 µm. F. Heatmaps from RNA-seq analysis of Cth1 mutant sperm show downregulation of genes involved in protein autophosphorylation. G. Heatmaps from RNA-seq analysis of Cth1 mutant sperm show downregulation of genes involved in the electron transport chain. H. Immunostaining for the DNA damage markers RAD51 reveals elevated levels of DNA damage in Cth1 mutant testes compared to wildtype testes. Scale bar 100 µm. I. Prussian blue staining of testes sections reveals increased accumulation of iron-positive particles in Cth1 mutant testes compared to wild type. Scale bar 20 µm. J. Immunostaining for LC3B shows increased signal intensity in Cth1 mutant testes sections relative to controls, consistent with elevated autophagic activity. Negative control sections show no detectable staining. Scale bar 100 µm.

